# Bovine blood and milk T-cell subsets in distinct states of activation and differentiation during subclinical *Staphylococcus aureus* mastitis

**DOI:** 10.1101/2023.01.12.523729

**Authors:** Nisha Tucker, Patricia Cunha, Florence B. Gilbert, Marion Rambault, Kamila Reis Santos, Aude Remot, Pierre Germon, Pascal Rainard, Rodrigo Prado Martins

## Abstract

T-lymphocytes are key mediators of adaptive cellular immunity and knowledge about distinct subsets of these cells in healthy and infected mammary gland secretions remains limited. In this study, we used a multiplex cytometry panel to show that staphylococcal mastitis causes the activation of CD4^+^, CD8^+^ and γδ T-cells found in bovine milk. We also highlight remarkable differences in the proportions of naïve and memory T-cells subsets found in blood and milk. These observations will contribute to a better understanding of cell-mediated immune mechanisms in the udder and to the development of new therapeutic and preventive strategies targeting mastitis.

## 1. Introduction

T-lymphocytes (T-cells) are essential for the establishment and maintenance of cell-mediated immunity (CMI). Upon recognition of a specific antigen, T-cells proliferate and acquire effector functions that limit pathogen dissemination. These cells subsequently contribute to protection of the host from re-infection by insuring the maintenance of long-lived memory cells. Although ruminant T-cell biology has long been a topic of interest, the scarcity of adapted reagents and methods has limited new discoveries.

Accumulating evidence highlights a major role of T-cells in the defence of the mammary gland (MG) against bacterial infections (mastitis). Mastitis represents the most frequent disease of dairy cows and the primary cause of antibiotics use in dairy farming, *Staphylococcus aureus* being the main cause of contagious and chronic infections in cows (Soltys and Quinn 1999). A broader description of T-lymphocyte populations found in MG tissue and secretions, under physiological and pathological conditions, has been pointed out as a necessity for the development of effective mastitis vaccines (Rainard *et al.* 2022).

We took advantage of a multiplex cytometry panel recently published by Roos *et al.* (2022) to identify fifteen distinct subsets of CD4^+^, CD8^+^ and γδ T-cells in the blood and milk of dairy cows carrying chronic subclinical mastitis by *S. aureus*.

## 2. Material and methods

### 2.1 Animals and sample collection

Blood and milk samples were collected from five Prim’Holstein cows in lactation bred in a farm located in the French department Marne-et-Loire. Studied animals were diagnosed with chronic subclinical mastitis caused by a common strain of *S. aureus* in at least one mammary quarter. Animal handling and blood sampling were conducted with the approval of Ethics Committee of Val de Loire (France, DGRI’s agreement APAFIS#23901-20200203095344) in strict accordance with all applicable provisions established by the European directive 2010/63/UE. Blood samples from the jugular vein were collected into 10 ml vacuum tubes containing EDTA (Vacutainer K2-EDTA, BD) and kept at room temperature until processing (< 3 h). For milk sampling, teats were washed, dried with paper towels and disinfected with alcohol-soaked swabs. The first four milk squirts were discarded then 500 ml of milk was collected in a sterile container and kept in a cooler box with frozen gel packs until processing (< 3 h).

### 2.2 Cell separations

Blood samples were processed for the separation of buffy coat cells as previously described (Cunha *et al.* 2022). For the separation of milk cells, somatic cells were initially counted using an automated cell counter (Fossomatic model 90; Foss Food Technology, Hillerod, Denmark). Milk was then diluted in DPBS (Gibco) in a 1:1 ratio and centrifuged for 30 min at 1400 g at 20 °C. The fat layer and supernatant were removed and cell pellets were washed with 10 ml of FACS buffer (DPBS without Ca2^+^ and Mg2^+^ supplemented with 2% (v/v) normal goat serum and 2mM EDTA) and resuspended in the same buffer for further analysis.

### 2.3 Microbiological analysis of milk samples

Fifty microliters of milk were streaked on blood agar plates (Merck Millipore) and incubated at 37 °C for 48h. The identification of haemolytic colonies positive in Gram, catalase and clumping factor/coagulase tests was verified by PCR using *S. aureus* specific primers as described in Gilbert *et al.* (2006).

### 2.4 Immunophenotyping by flow cytometry

Surface markers were labelled using the antibodies and conditions described in **Supplementary Table 1.** Following antibody incubations, dead cells were stained using the Fixable Viability Dye eFluor 450 (eBioscience). Cells were examined by flow cytometry using a BD LSR Fortessa^™^ X-20 and data were analysed with the Kaluza software (Beckman Coulter). Dead cells were excluded from the analysis and the strategy described by Roos *et al.* (2022) was employed to study fifteen lymphocyte subpopulations, as shown in **Figure 1A** and **Supplementary Figure 1**.

**Figure 1.**
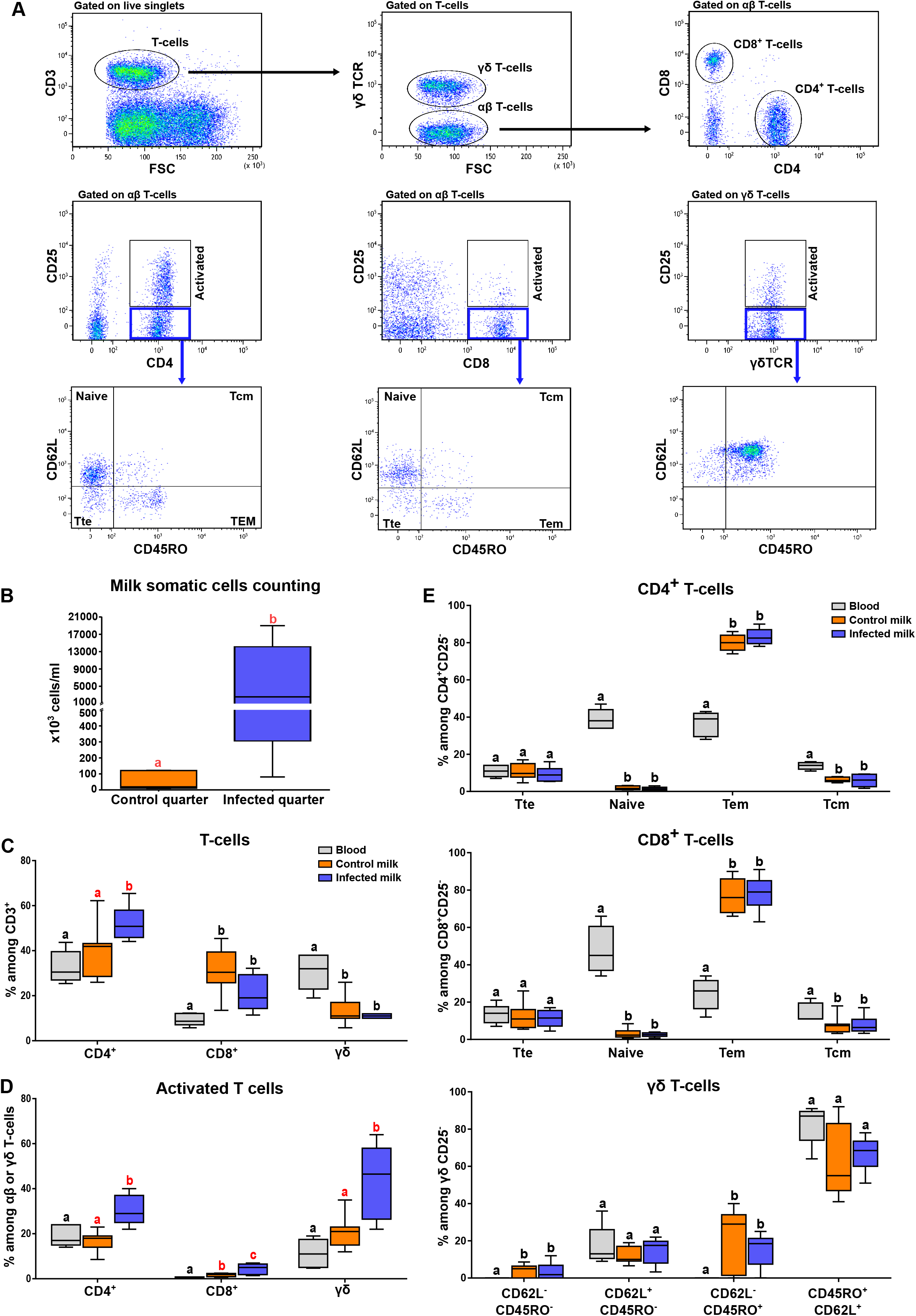
T-cell populations found in bovine blood and milk during staphylococcal mastitis. **A.** Gating strategy set up to analyse distinct CD3^+^ lymphocyte populations by flow cytometry. TTE, Tem and Tcm denote T terminal effector, T effector memory and T central memory populations, respectively. Results of a representative analysis with blood cells are shown. **B.** Somatic cells counting on milk from healthy (control) and *S. aureus*-infected mammary quarters. **C.** Percentage of CD4^+^, CD8^+^ and γδ T-cells among CD3^+^ cells. **D.** Percentage of activated (CD25^+^) among CD4^+^, CD8^+^ or γδ T-cells. **E.** Percentage of naive, effector and memory T-cell subpopulations among non activated (CD25^-^) CD4^+^ (upper graph), CD8^+^ (middle graph) or γδ T-cells (lower graph). In all plots, data were analysed using the Mann-Whitney test. Different letters indicate significant statistical difference (p<0.05) among blood, control and infected milk. Significant differences between control and mastitic milk are highlighted in red.

### 2.5 Statistical analysis

Data were analysed using R-Studio (version 2022.07.2+576) and plotted using GraphPad Prism, Version 6.0 (GraphPad Software Inc.). Pairwise comparisons were carried out using the Mann-Whitney test. For the box-plots, statistically significant differences (p<0.05) between groups are represented by different letters.

## Results and discussion

Microbiological analyses confirmed the presence of *S. aureus* in six mammary quarters and seven quarters in which microbiological growth was absent were used as controls. The occurrence of subclinical mastitis was further confirmed by the observation of somatic cell counts (SCC) above 200.10^3^ cells^ml^ in milk samples positive for *S. aureus,* whereas SCC in control quarters were below this threshold (**Figure 1B**). Next, we set out to explore the T-cell populations found in the blood as well as in milk from affected and control quarters. In agreement with previous reports (Soltys and Quinn 1999, Rainard *et al.* 2022), CD8^+^ T-cells were shown to be more abundant in milk than in blood and mastitis by *S. aureus* led to a decrease of these cells concurrent with an increase of CD4^+^ T-cells. Additionally, subclinical mastitis led to no significant changes in the percentage of γδ T-cells in milk, these cells being more frequently observed in blood (**Figure 1C**). Previous evidence demonstrate that γδ T-cells, reported as a major regulatory T-cell subset in the bovine species (Guzman *et al.* 2014), are more abundant in milk during streptococcal and staphylococcal clinical mastitis (Soltys and Quinn 1999) and might contribute to the resolution of acute inflammation in the MG (Slama *et al.* 2021). The CD25 molecule corresponds to the α chain of the IL-2 receptor and its expression has been associated with the functional activation of conventional and regulatory T-cells (Zelenay *et al.* 2005). As depicted in **Figure 1D**, milk from infected quarters contained a higher percentage of activated γδ T-cells and this indicates that although these lymphocytes are not more recruited to the MG and milk during the chronic phase of mastitis, they might exert a regulatory function in this context.

Activated CD4^+^ T-cells were also more frequently found in infected milk (**Figure 1D**) in agreement with the reported role of these cells as local regulators of IL17 and IFNγ responses in a murine model of mastitis by *S. aureus* (Zhao *et al.* 2015) and during the antigen-specific immune responses in the bovine MG (Rainard *et al.* 2022). Costimulatory signals driven by CD4^+^ T cells are also crucial for the differentiation of CD8^+^ T cells, the expression of CD25 on pathogen-specific CD8^+^ T cells being dependent on CD4^+^ T cell help (Obar *et al.* 2010). Interestingly, CD25-expressing CD8^+^ T cells were more abundant in milk, even in non-infected quarters, than in blood (**Figure 1D**). In agreement with observations in breast milk (Wirt *et al.* 1992, Sabbaj *et al.* 2005), we found out a predominance of T effector memory cells (Tem) and low proportions of T central memory cells (Tcm) and naïve T-cells in both control and infected milk when compared to blood (**Figure 1E**). Mastitis by *S. aureus* often result in persistent infections in bovine and affected animals do not develop protection against re-infection. This particularity has been associated with the host’s inability to generate effective local memory responses, in agreement with our results showing that activated but not memory T-cells are more abundant in infected milk. Activated CD8^+^ T cells from dry MG secretions have been previously reported to supress the proliferative response of CD4^+^ lymphocytes during bovine mastitis by *S. aureus* (Park et al. 1993). It is therefore tempting to infer that during staphylococcal mastitis, activated CD4^+^ T-cells could cause the activation of regulatory CD8^+^ lymphocytes, which in turn hamper T-cell clonal expansion and differentiation into long-lived, *S. aureus*-specific stable memory T cells. Nevertheless, further research is necessary to address this hypothesis.

Surprisingly, γδ T-cells were shown to be predominantly CD45RO^+^CD62L^+^ in milk and blood, a CD45RO^+^CD62L^-^ population being observed only in milk (**Figure 1E**). Since memory γδ T-cells have been associated with effective protection to *S. aureus* in mice (Lalor and McLoughlin 2016), the role of these cells in cattle deserves to be clarified.

These observations will contribute to a better understanding of T-cell populations found in milk in physiological and pathological conditions and might pave the way for further studies focused on clarifying how T-cells modulate immune response in the MG.

## Acknowledgements

The authors thank Yves Le Vern for his skilful support with the flow cytometry analysis.

## Contributions

Conceptualization, RPM; Experimental work, NT, PC, FBG, KRS and RPM; Data analysis, RPM, NT; Resources, MR, AR, and PG; Writing-Original Draft Preparation, RPM; Writing-Review & Editing, RPM, PR, PG, FBG and NT; Supervision, RPM and PC; Funding Acquisition, RPM.

---

**Supplementary Table 1.**
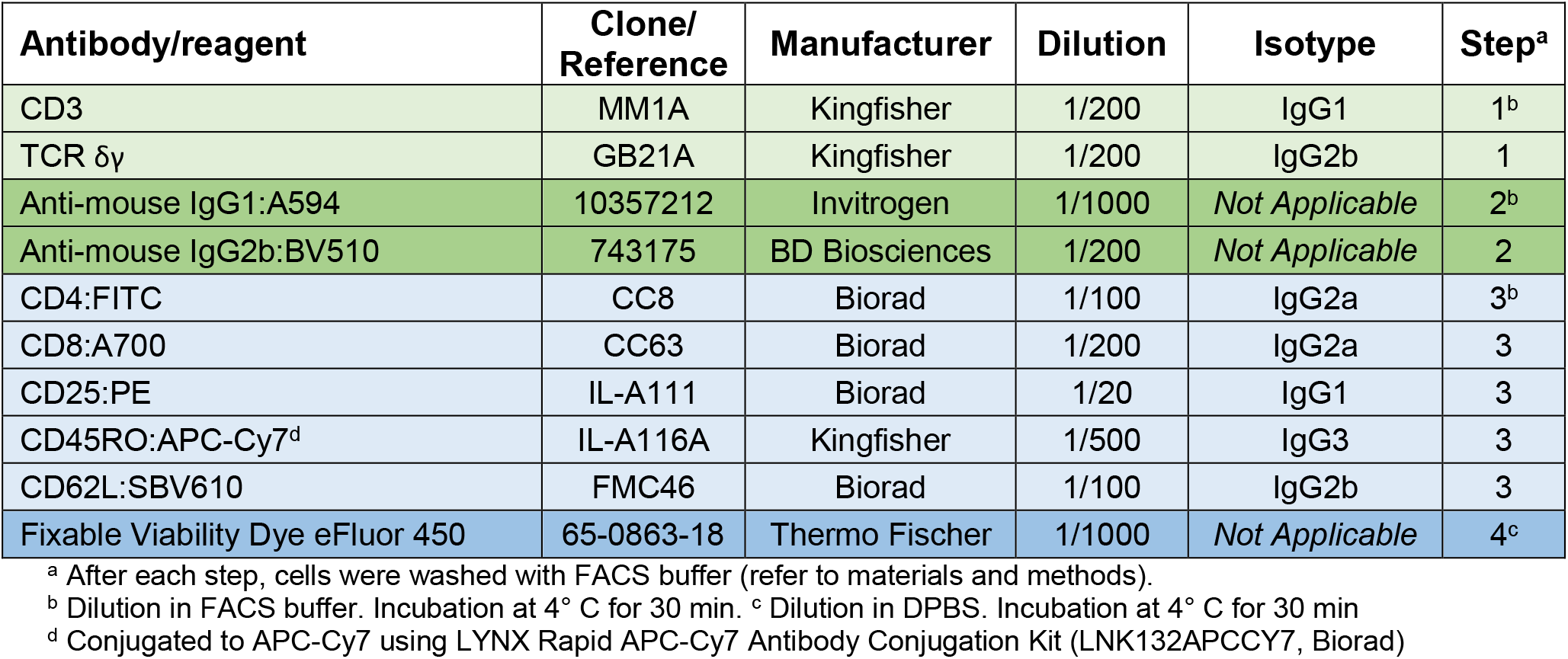
Antibodies and conditions used for the phenotyping of bovine lymphocyte populations by flow cytometry

