## Supplementary Figure 1 for "Bovine blood and milk T-cell subsets in distinct states of activation and differentiation during subclinical *Staphylococcus aureus* mastitis"

**Supplementary Figure 1 -** Detailed gating strategy employed for the analysis of milk and blood T-lymphocytes subsets. TTE, Tem and Tcm denote T terminal effector, T effector memory and T central memory populations, respectively. Results of a representative analysis with blood cells are shown. FVD means Fixable Viability Dye

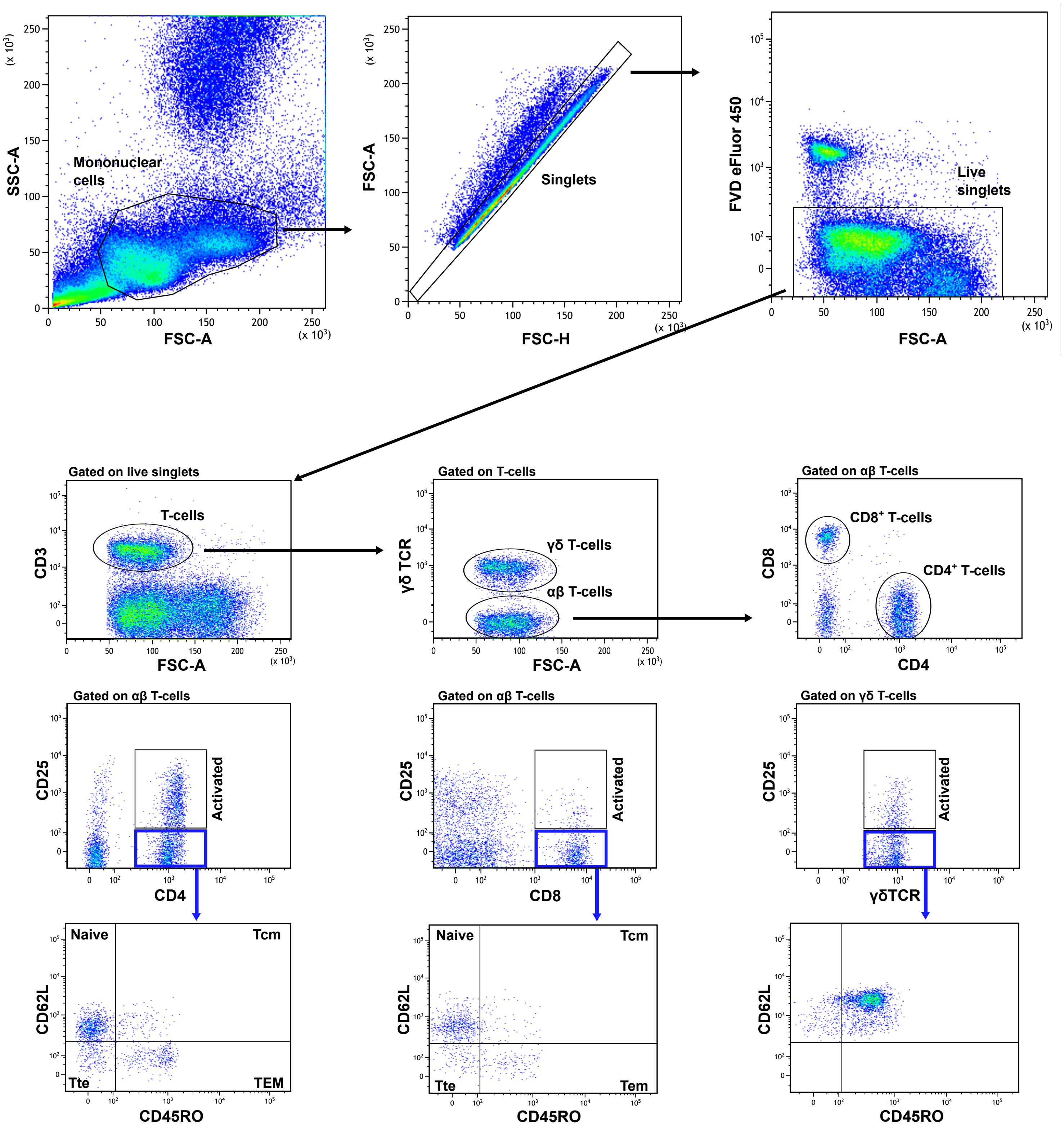
