## Supplementary table 1 for "Bovine blood and milk T-cell subsets in distinct states of activation and differentiation during subclinical *Staphylococcus aureus* mastitis"

**Supplementary Table 1 –** Antibodies and conditions used for the phenotyping of bovine lymphocyte populations by flow cytometry

| **Antibody/reagent** | **Clone/ Reference** | **Manufacturer** | **Dilution** | **Isotype** | **Step[a]** |
|---|---|---|---|---|---|
| CD3 | MM1A | Kingfisher | 1/200 | IgG1 | 1[b] |
| TCR δγ | GB21A | Kingfisher | 1/200 | IgG2b | 1 |
| Anti-mouse IgG1:A594 | 10357212 | Invitrogen | 1/1000 | *Not Applicable* | 2[b] |
| Anti-mouse IgG2b:BV510 | 743175 | BD Biosciences | 1/200 | *Not Applicable* | 2 |
| CD4:FITC | CC8 | Biorad | 1/100 | IgG2a | 3[b] |
| CD8:A700 | CC63 | Biorad | 1/200 | IgG2a | 3 |
| CD25:PE | IL-A111 | Biorad | 1/20 | IgG1 | 3 |
| CD45RO:APC-Cy7[d] | IL-A116A | Kingfisher | 1/500 | IgG3 | 3 |
| CD62L:SBV610 | FMC46 | Biorad | 1/100 | IgG2b | 3 |
| Fixable Viability Dye eFluor 450 | 65-0863-18 | Thermo Fischer | 1/1000 | *Not Applicable* | 4[c] |

[a] After each step, cells were washed with FACS buffer (refer to materials and methods).
[b] Dilution in FACS buffer. Incubation at 4° C for 30 min. [c] Dilution in DPBS. Incubation at 4° C for 30 min
[d] Conjugated to APC-Cy7 using LYNX Rapid APC-Cy7 Antibody Conjugation Kit (LNK132APCCY7, Biorad)
